# Resolving Individual-Atom of Protein Complex using Commonly Available 300-kV Cryo-electron Microscopes

**DOI:** 10.1101/2020.08.19.256909

**Authors:** Kaiming Zhang, Grigore D. Pintilie, Shanshan Li, Michael F. Schmid, Wah Chiu

**Affiliations:** Department of Bioengineering and James H. Clark Center, Stanford University, Stanford, CA 94305, USA; Division of CryoEM and Bioimaging, SSRL, SLAC National Accelerator Laboratory, Menlo Park, CA 94025, USA

## Abstract

Breakthroughs in single-particle cryo-electron microscopy (cryo-EM) technology have made near-atomic resolution structure determination possible. Here, we report a ∼1.35-Å structure of apoferritin reconstructed from images recorded on a Gatan K3 or a Thermo Fisher Falcon 4 detector in a commonly available 300-kV Titan Krios microscope (G3i) equipped with or without a Gatan post-column energy filter. Our results demonstrate that the atomic-resolution structure determination can be achieved by single-particle cryo-EM with a fraction of a day of automated data collection. These structures resolve unambiguously each heavy atom (C, N, O, and S) in the amino acid side chains with an indication of hydrogen atoms’ presence and position, as well as the unambiguous existence of multiple rotameric configurations for some residues. We also develop a statistical and chemical based protocol to assess the positions of the water molecules directly from the cryo-EM map. In addition, we have introduced a B’ factor equivalent to the conventional B factor traditionally used by crystallography to annotate the atomic resolution model for determined structures. Our findings will be of immense interest among protein and medicinal scientists engaging in both basic and translational research.

## Introduction

X-ray crystallography and single particle cryo-electron microscopy (cryo-EM) have become wide-spread, routine tools for characterizing biochemically purified specimens. cryo-EM has resolved over four thousand structures at near atomic resolutions (2-4 Å^1^). It is rapidly becoming the method of choice for structure determination of membrane proteins, large assemblies, and multi-protein complexes partly because it does not require a crystal and partly because it can work with specimens with heterogeneous composition and/or conformation^2^, This powerful technique is now capable of resolving biological specimens to better than 2 Å resolution^3^ and has been used to solve 3.7-Å resolution structures of specimens as small as ∼40 kilodaltons^4^. Cryo-EM instrumentation is being installed in many academic and industrial institutions worldwide. In order to make this imaging modality to generate detailed atomic level structure in the classic chemistry context or the investigation space for drug-design development, one really needs to reach a structure resolution at least 1.5 Å or better.

In our study, we achieved cryo-EM maps of apoferritin, reconstructed from the images collected from the commonly available 300-kV Titan Krios microscopes at 1.34 Å using K3 detector and at 1.36 Å using Falcon4 detector, showing the detailed structural information including the well-resolved amino acid side chain atoms with an indication of hydrogen atoms’ presence and position, and the existence of multiple rotameric configurations for some residues.

Since the resolution used in cryo-EM is neither a conventional optics criterion nor the detection of diffraction spots in a diffraction pattern of a crystal. Evaluation of the high resolution structure details in the cryo-EM maps and their quantitative assessments are necessary and important for the field. We recently developed a parameter called Q-score^3^ to assess the high-resolution cryo-EM maps up to 1.5 Å. In this study, we further updated the empirical formulation of Q-score for its usability for structure data up to 1 Å. We also develop a protocol to assess the position of the water molecules solely from the cryo-EM map as well as to annotate the atomic model to be reflective of the actually observed density which could be resolved differently throughout the map.

## Results and Discussion

Two years ago, we obtained a 1.75-Å apoferritin structure^3^ using ∼70,000 particle images through 10-h of data collection using a Gatan K2 detector in a Titan Krios G3i electron microscope. Recently the K3 and Falcon4 detectors were installed on two of our Titan Krios G3i electron microscopes at the Stanford-SLAC Cryo-EM Center, providing us with three times the data collection rate. In this study, using the same apoferritin, we first collected a new dataset using the K3 detector to find out what is the highest resolution structure achievable. The dataset was collected in the electron counting mode using “Faster Acquisition’’ in EPU, with a throughput of ∼520 movie stacks per hour and a data acquisition result of 8,000 movie stacks per 16 h (**Table S1**). Using a standard image processing pipeline^5^ (**Fig. S1a**), a density map of apoferritin with a resolution of 1.34-Å was obtained from ∼900,000 particles (**Fig. 1a-d**). In addition, we also collected a second dataset using a Falcon4 detector in the electron counting mode (mrc format, not EER format) with the throughput of ∼500 movie stacks per hour, resulting in ∼ 7,700 movie stacks from a 15-h data collection. A 1.36-Å resolution map of apoferritin was obtained from ∼ 500,000 particles (**Fig. S1b-e, Table S1**). Both maps have comparable resolution based on the Fourier Shell correlation of 0.143 threshold^6^ and similar cumulative B-factor as estimated from reconstructions with varying numbers of particles^7^ (**Fig. S1f-g**).

**Fig 1.**
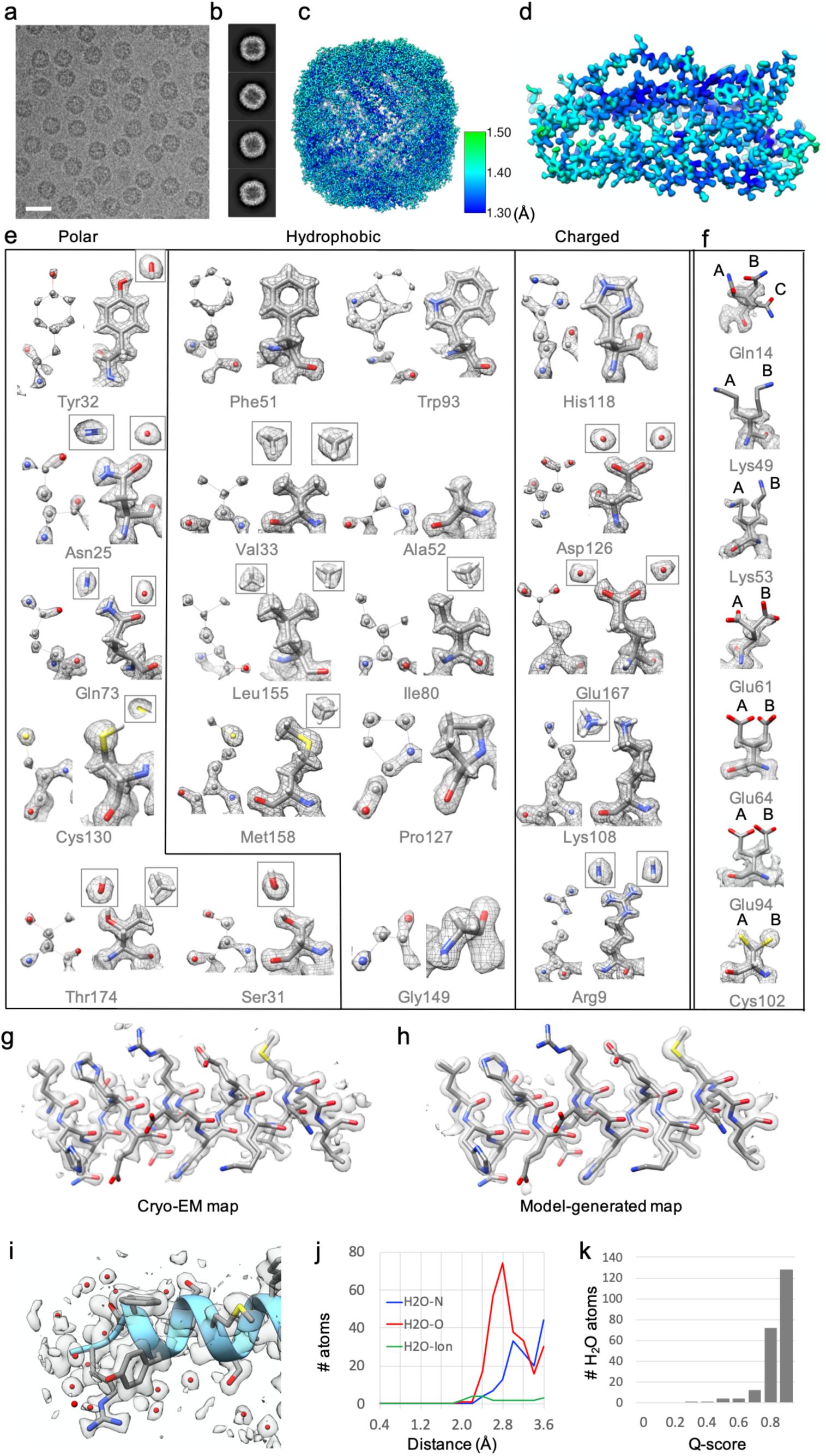
Atomic resolution structure of apoferritin determined from a commonly used 300kV Titan Krios electron microscope with K3 detector. **a**. Representative motion-corrected cryo-EM micrographs. The scale bar represents 200 Å. **b**. Reference-free 2D class averages of computationally extracted particles. **c**. Resolution variation maps for the final 3D reconstruction. **d**. Cryo-EM density map of an extracted single subunit. **e**. Twenty representative amino acids extracted from the 1.34-Å map. The amino acids were selected based on the type of side chain (polar, charged, and hydrophobic). Each residue is shown on a higher contour level (left in every box) or lower contour level (right in every box), showing separable/resolved atoms or shapes of atoms including hydrogen atoms, respectively. **f**. Representative residues with alternate conformations of side chains. A/B/C represents different side chain conformations. The residues in e and f are shown by element (grey, carbon; red, oxygen; blue, nitrogen; yellow, sulphur; white, hydrogen). **g, h**. A representative helix was extracted from the cryo-EM density map (g) and the model-generated map using B’-factors (h). **i**. Water molecules are shown around a small portion of the helix. **j**. A radial distance plot between water and O atoms in the protein shows a sharp peak at 2.8 Å. **k**. A histogram of Q-scores for placed waters shows that most are placed in well-resolved peaks with Q-scores of 0.8 and higher.

Our maps show the individual atom density at properly chosen contour level, all well-resolved side-chain atom densities, and even some indications of hydrogen densities as exemplified from the 1.34-Å map (**Fig. 1e**). The presence and direction of the density for hydrogen, even though it is not distinctly resolved, is visually compelling. Furthermore, the difference between the oxygen and the nitrogen in Asn and Gln, where the N has extended density for its H atoms, whereas the O is rounded, allows one to distinguish these two atoms, which informs whether the atom is a hydrogen bond donor or acceptor. The same is the case for the O vs C in Thr, wherein the terminal methyl group is roughly triangular in shape, and the O again is rounder, and allows one to unambiguously differentiate Thr from Val. However, the side chains of some residues cannot be clearly identified due to multiple rotameric conformations (**Fig. 1f**), which may be caused by their inherent or electron radiation-induced dynamic properties.

The Molprobity and PDB reports of our two models are ranked very highly in all the assessment scores on the adherence of models to the chemical properties of proteins (**Fig. S2**). Generally speaking, it is difficult to assess the fit of a model to a density map by visual display, partly because of the choice of contour and partly because of the variation of resolvability throughout the map (**Fig. 1c-d, Fig. S1e**). Though the overall resolutions of maps are reported based on the Fourier shell correlation (**Fig. S1f**), some side chains and residues are less resolved. We used Q-scores, a recently proposed cryo-EM structure validation method, to measure the resolvability of individual atoms^3^. Q-scores were shown to correlate strongly to the resolution of the map with the best score normalized to 1.0. **Fig. 2a** shows a per-residue Q-score plot for our two maps to range between 0.85 and 0.88; most residues have average Q-scores at or above the expected level for this resolution^3^. However, a few dips in the plot can be seen, indicating lower average Q-scores at turns or loops between helices. Visual inspection of the residues with lower Q scores confirms that the side chains are less resolved for these residues. Such residues tend to have alternate conformers (**Fig. 1f**) and be on the exterior surface of the complex.

**Figure 2.**
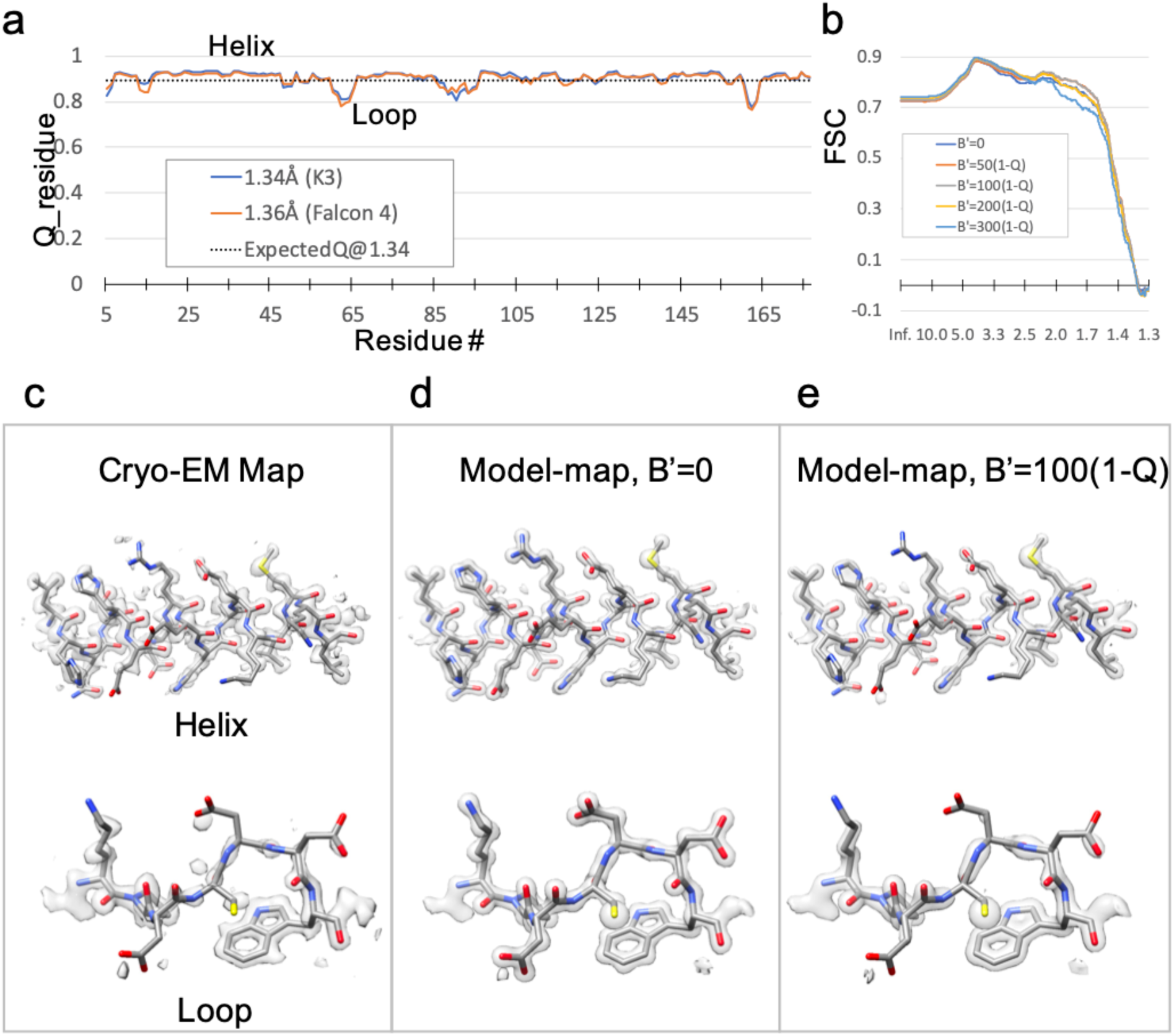
Q-scores and B’ factors. **a**. Plots of per-residue Q-score for 1.34 Å and 1.36 Å maps. Most residues have Q-scores above the expected Q-score at a resolution of 1.34 Å, with just a few dips which occur mostly in loop regions. **b**. FSC plots between the 1.34 Å map and model are shown, using several B’ scaling factors (see Methods). The scaling factor of B’=100(1-Q) has the highest FSC correlations at all resolutions. **c-e**. Two extracted helical and loop regions to show the consistency between cryo-EM map and model-derived map. In (c), the cryo-EM map is shown. In (d), a model-generated map with B’-factors of 0 is shown; all atoms are inside the contour shown, unlike in the cryo-EM map. In (e), a model-generated map with B’-factors calculated from Q-scores is shown; here, atoms that have lower resolvability in the cryo-EM map are similarly un-resolved in the model-generated map.

Traditionally, B-factor is used to assess the atom position uncertainty in crystallography and is a weighting factor to allow computing a model-based map identical to the experimental map. Note that the crystallographic B-factor is estimated by iterative and simultaneous refinements among diffraction spot indexing, Fourier amplitudes between observed and computed values, and Fourier phase estimation. However, there is currently no routine procedure to derive an equivalent per-atom B-factors in cryo-EM modeling. We thus introduce B’ factors derived from per-atom Q-scores; the calculation involves a single scaling factor; the optimum scaling factor is determined empirically by testing which value makes the resulting model-map matched the cryo-EM map better by FSC (**Fig. 2**). The B’ factor, which will be deposited to the PDB, serves the same purpose as the crystallographic B-factor in such a way that we can compute a model-based map that can match well with the experimental cryo-EM density map (**Fig. 1g-h, Fig. 2, and Methods**).

Resolving water molecules is an important index for assessing the quality of a true atomic resolution map. We assigned water molecules in our maps using a new procedure based on three criteria: a signal to noise threshold to ignore background noise (2-sigma/RMSD above average), the distance between putative water to the closest protein atoms, and the rules that distinguish water and ions as outlined in ref^8^ (**Fig. 1i-k; Fig. 3a-d**). The distributions show that the procedure places water on well-resolved peaks with Q-scores of 0.5 and higher, even though Q-scores were not used in the procedure itself. The radial-distance plot shown in **Fig. 1j** shows a peak at 2.8 Å between water atoms to nearby O atoms in protein, as expected. In comparison with other apoferritin maps determined at better than 2 Å, more water molecules were found in higher-resolution maps: 217 waters in the 1.25 Å map, 170 waters in our 1.34 Å, 165 waters in our 1.36 Å map, and 126 waters in our previous 1.75 Å map (per protomer) (**Fig. 3e-f**). Sharper peaks were observed at higher resolution, meaning that higher resolution maps localize water molecules more accurately. Molprobity results showed only a few of the waters placed with our procedure were found to clash (3 in the 1.34 Å map and 8 in the 1.36 Å map) (**Fig. S2**).

**Figure 3.**
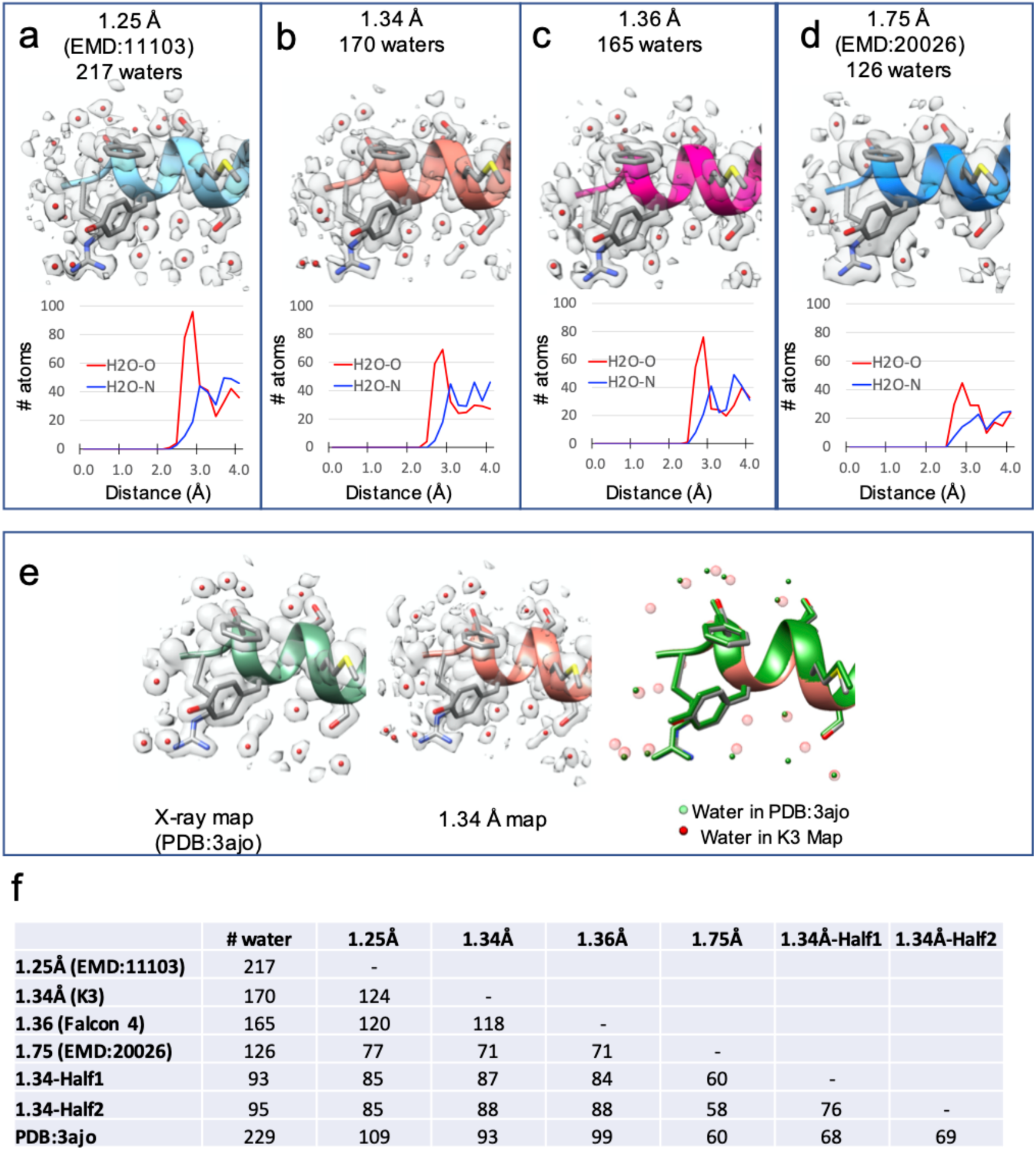
Water finding in apoferritin maps at different resolutions. **a-d**. Extracted regions from four cryo-EM maps show water molecules near protein atoms, along with radial-distance plots. The latter shows peaks at a distance of 2.8 Å from protein O atoms. **e**. Comparison of water molecules between an X-ray structure (PDB: 3ajo) and our 1.34 Å map. **f**. Comparison of water molecules placed in different cryo-EM maps. The first column shows the number of water molecules found in each map. The other columns are an N ⨯N comparison of the number of waters that are within 1 Å of each other in two different maps.

We compared water molecules placed in different cryo-EM maps (1.25 Å [EMD:11103], 1.34 Å, 1.36 Å, 1.75 Å [EMD:20026], and two half-maps of the 1.34 Å structure) and an X-ray map [pdb:3ajo]. When comparing different cryo-EM maps, an average of 76% water molecules found in one map were also found within 1.0 Å in the other maps (**Fig. 3e-f**). Based on this comparison, it is encouraging to note that a high percentage of water molecules placed in cryo-EM maps agree, suggesting good reproducibility of water positions across data sets recorded in different electron microscopes and sample preparations. When comparing cryo-EM maps to the X-ray structure, a lower percentage (36%) of water molecules were within the same 1 Å distance. Given the differences in sample preparation (like concentration and buffer) or conditions (crystal packing of X-ray model vs in-solution state in cryo-EM), it is not surprising that there are differences in water positions. Such discrepancy also applies to ion placements. In placed ions, positions compared amongst cryo-EM maps showed lower similarity (34%). This lower similarity could be due to higher sensitivity of ion locations due to different sample conditions. Water and ion identification is still under active research even in atomic resolution X-ray structure^8^, and it is an emerging but potentially important area in cryo-EM map analysis.

Recently, two unpublished preprints report 1.22 and 1.25-Å-resolution apoferritin structures^9,10^ using new electron optics (i.e. cold field emission gun, second-generation spherical aberration lens corrector/monochromator), which are aimed to minimize the deleterious effects of the energy spread to reduce the high-resolution signals^11^. In our study, we show two ∼1.35-Å structures of the apoferritin without these new hardware upgrades. We further selected four representative residue types for a detailed comparison at the individual atom level among these cryo-EM maps, a 1.01-Å crystal structure and a 1.75 Å cryo-EM map (**Fig. 4**). As expected, the two cryo-EM maps better than 1.3 Å have slightly higher Q-scores and atomic separation was slightly better. From the overall and detailed evaluation, our 1.34 and 1.36 Å cryo-EM maps show virtually the same features as the 1.22 Å and 1.25 Å cryo-EM maps and the 1.01 Å crystal structure, and all are better than the 1.75 Å map. Our results demonstrate that an atomic resolution structure can be obtained on a commonly available 300 kV microscope using the K3 or Falcon4 detectors with or without an energy filter (**Table S1**), which thus gives the current majority of users the possibility of performing single-particle cryo-EM analysis at the atomic resolution level.

**Figure 4.**
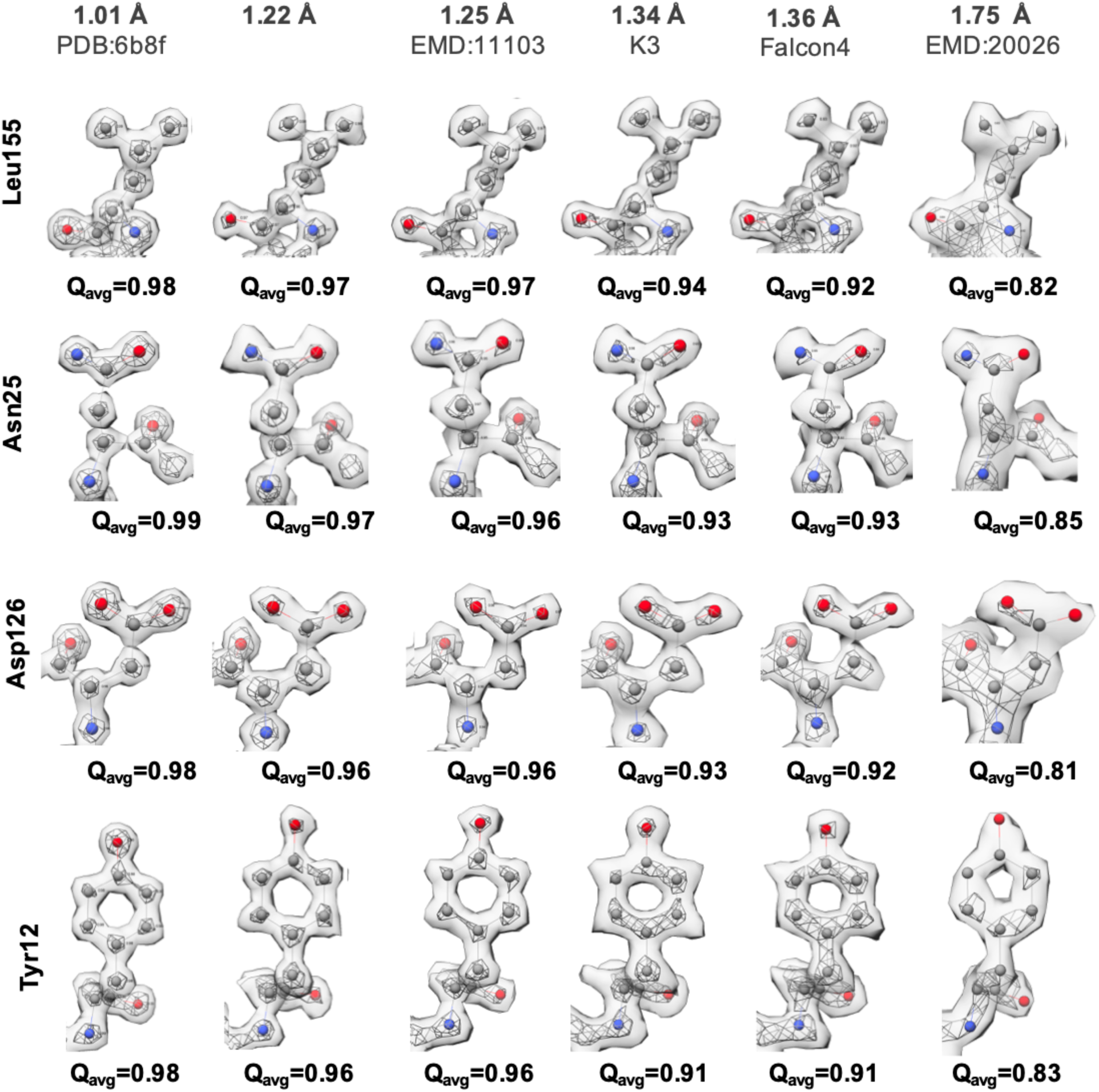
Comparing the resolvability of apoferritin structures at multiple resolutions by Q-score. Four different residues were selected based on the type of side chain (polar, charged, and hydrophobic). The residues are shown by element (grey, carbon; red, oxygen; blue, nitrogen). The 1.22 Å map was downloaded from Scheres lab (ftp://ftp.mrc-lmb.cam.ac.uk/pub/scheres/atomic/).

Finally, we would like to remind that apoferritin, with its high stability, rigidity, and symmetry, can easily be resolved at atomic resolution by either cryo-EM or X-ray crystallography. However, many biological samples are compositionally or conformationally heterogeneous, as well as often difficult to prepare them vitrified with the current cryo-freeze plunge method^12^. These technical hurdles can hinder solving their structures at high resolution. Achieving atomic resolution structures is not yet a routine task, but with the further development of cryo-specimen preparation, hardware, and software, it should be possible to apply this approach to an even broader spectrum of macromolecules in the context of chemistry to understand the mechanism and/or to apply it in the drug design pipeline.

## Materials and Methods

### Cryo-EM sample vitrification and data acquisition

Three microliters of the apoferritin samples with 1.5 mg/ml or 0.2 mg/ml were applied onto glow-discharged 200-mesh R2/1 Quantifoil grids (used for the K3 dataset) or 200-mesh R2/1 Quantifoil grids coated with continuous carbon film (Quantifoil, used for the Falcon4 dataset), respectively. The grids were blotted for 4 s or 2 s and rapidly cryocooled in liquid ethane using a Vitrobot Mark IV (Thermo Fisher Scientific) at 4°C and 100% humidity. The first dataset (K3 dataset) was imaged in a Titan Krios cryo-electron microscope (Thermo Fisher Scientific) operated at 300 kV with GIF energy filter (Gatan) at a magnification of 215,000× (corresponding to a calibrated sampling of 0.4 Å per pixel). Micrographs were recorded by EPU software (Thermo Fisher Scientific) with a Gatan K3 Summit direct electron detector, where each image was composed of 30 individual frames with an exposure time of 0.5 s and an exposure rate of 90 electrons per second per Å^2^. A total of 8,034 movie stacks were collected. The second dataset (Falcon4 dataset) was imaged in another Titan Krios cryo-electron microscope (Thermo Fisher Scientific) operated at 300 kV at a magnification of 155,000× (corresponding to a calibrated sampling of 0.502 Å per pixel). Micrographs were recorded by EPU software (Thermo Fisher Scientific) with a Falcon4 direct electron detector, where each image was composed of 40 individual frames with an exposure time of 2 s and an exposure rate of 20 electrons per second per Å^2^. A total of 7,734 movie stacks were collected.

### Single-particle image processing and 3D reconstruction

All micrographs were first imported into Relion for image processing. The motion-correction was performed using RELION’s own implementation and the contrast transfer function (CTF) was determined using CTFFIND4^13^. For the K3 dataset, 6,951 micrographs were selected with a defocus range from −0.35 - −1.3 μm, and “rlnCtfMaxResolution < 4.5”. For the Falcon4 dataset, 5,427 micrographs were selected with a defocus range from −0.3 to −1.3 μm, and “rlnCtfMaxResolution < 4”. All particles were autopicked using the NeuralNet option in EMAN2^14^. Then, particle coordinates were imported to Relion, where the poor 2D class averages were removed by two rounds of 2D classification. The initial models for both datasets were built in cryoSPARC^15^ using the ab-initio reconstruction option with Octahedral symmetry applied. For the K3 dataset, 1,176,336 particles were picked and 902,455 were selected after 2D classification. For the Falcon4 dataset, 707,350 particles were picked and 500,643 were selected after 2D classification. The 3D refinement was performed using the particle images selected from 2D classification with further “CTF refinement and Bayesian polishing” in Relion, a 1.34-Å map from the K3 dataset or 1.36-Å map from the Falcon4 dataset was obtained (Fig S1). Resolution for the final maps was estimated with the 0.143 criterion of the Fourier shell correlation curve. The figures were prepared using UCSF Chimera^16^.

### Q-score adjustment

Q-scores are calculated by correlating map values around each atom to a ‘reference Gaussian’. Previously, the width (sigma) of the reference Gaussian was set to 0.6 Å, which resulted in Q-scores of ∼1 at resolutions of 1.5 Å. With this sigma, Q-scores start to drop at higher resolutions. Hence, here we adjust sigma to 0.4 Å, so that Q-scores are now highest at a resolution of ∼1.2 Å, and a linear correlation can again be seen between Q-scores and resolution (**Fig. S3**).

### B’ calculations

We calculate B’ factors from atom Q-score using the following formula:

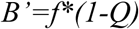

This formulation is such that atoms with higher Q-scores produce lower B’-factors, as they are better resolved. We determined the best scaling factor, f, by trying several values (0, 50, 100, 200, 300, 400), and observing which value caused the largest increase in the FSC.

### Segmentation-guided Water and Ion Modeling (SWIM)

In this procedure, the cryo-EM map is first segmented using the watershed method, which produces regions corresponding to peaks in the map; the boundaries between regions are the lowest values in the map around these peaks. A threshold of 2-sigma above the mean density value in the map is used, and only voxels with density values above this threshold are included in the segmentation. The resulting regions are then sorted by volume (number of voxels in the region), and considered in decreasing order as follows:

1. For each region, take the point in it with the highest map value as its position (P).
2. For each nearby atom to P:
  a. If the atom is non-polar and non-charged (e.g. carbon atom) and is within 2.6 Å of P, P is ignored and the search continues with the next regions.
  b. If the atom is charged (e.g. O in GLU/ASP or N in LYS/ARG/HIS sidechains) and is within a distance of 1.9 to 2.4 to P, it is added to ChargedAtoms list.
  c. If the atom is polar (e.g. O or N and any other side chain):
    i. If the atom is within 1.9 to 2.5 Å to P, it is added to PolarAtoms list.
    ii. If the atom is within 2.5 to 3.3 Å to P, it is added to WaterAtoms list.
  d. If the ChargedAtoms list is not empty, P is added as a 2+ ion (e.g. Mg)
  e. Otherwise, if the PolarAtoms list is not empty, P is added as 1+ ion (e.g. Na)
  f. Otherwise, if the WaterAToms list is not empty, P is added as a water.

In the above process, regions are considered in order of size, from largest to smallest. Hence, because waters would be placed on a larger region near an O or N atom rather than a smaller one at a similar distance, they would otherwise clash with the water placed on the larger region first.

## Acknowledgment

We thank Drs. Fei Sun and Xiaojun Huang (Institute of Biophysics, CAS) for providing the apoferritin sample. We thank Drs. Corey Hecksel and Patrick Mitchell for expert maintenance of Stanford-SLAC Cryo-EM Center and the SLAC National Accelerator Laboratory for supporting the conduct of these studies during the COVID-19 pandemic shutdown. This work was supported by the National Institutes of Health grants (U24GM129564, R01GM079429, P41GM103832).

## Author contributions

K.Z. and W.C. conceived the study and designed experiments. K.Z. performed cryo-EM sample preparation, data collection, image processing, and map reconstruction. G.D.P. developed the model and map correlation parameters. K.Z., G.D.P. and S.L. prepared the figures. All analyzed the data and wrote the manuscript.

## Data deposition

Cryo-EM maps of the apoferritin structure in this study with their associated atomic models have been deposited in the wwPDB OneDep System under EMD accession code XXX and PDB ID code XXX.

---

**Figure S1.**
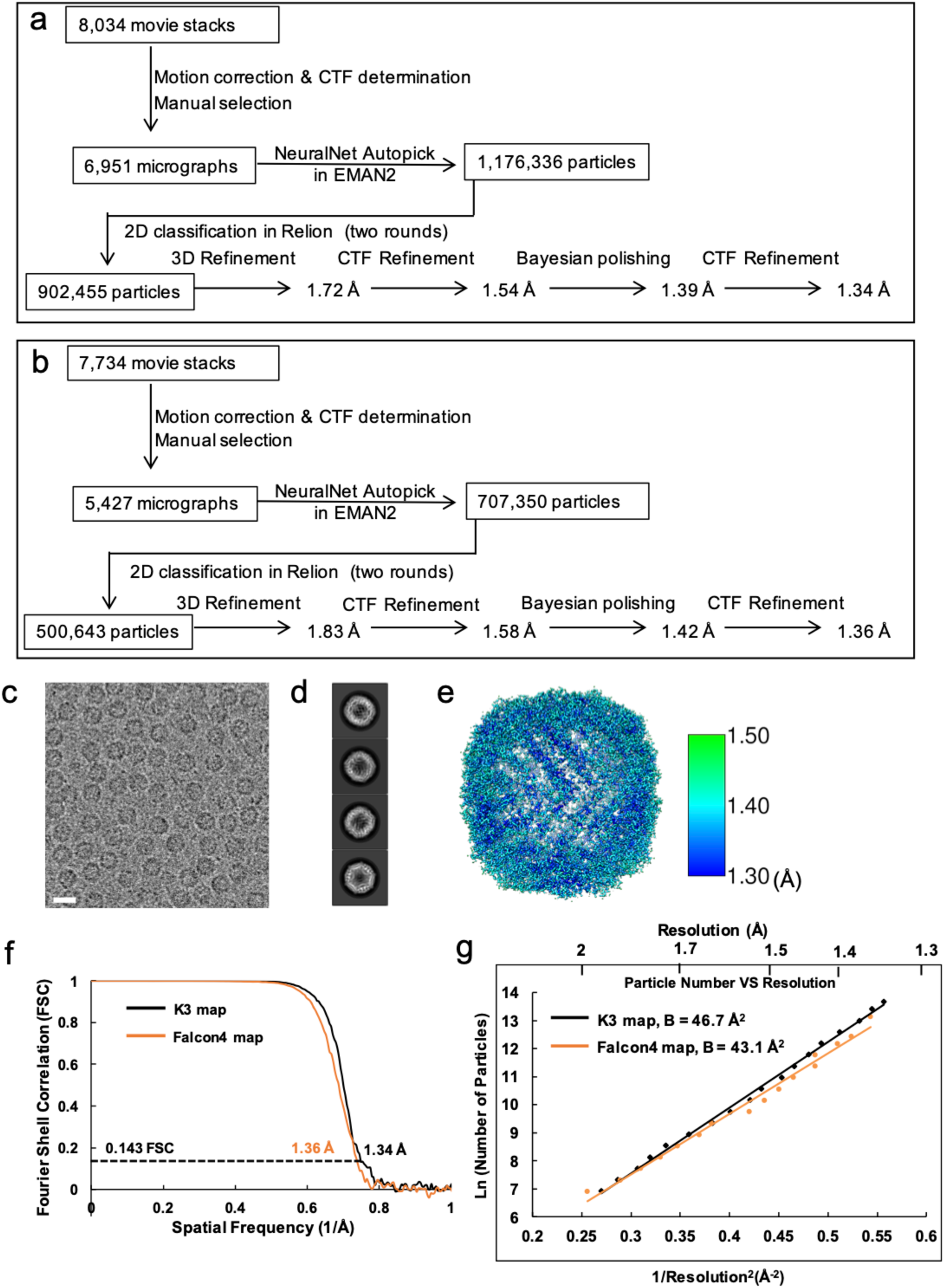
Single-particle cryo-EM analysis of apoferritin structures at atomic resolution from two datasets collected on K3 and Falcon4 detectors. **a**. Image processing workflow of the dataset collected on K3 detector. **b-e**. Data from Falcon4 dataset. **b**. Image processing workflow of the dataset collected on Falcon4 detector. **c**. Representative motion-corrected cryo-EM micrograph. The scale bar represents 200 Å. **d**. Reference-free 2D class averages of computationally extracted particles. **e**. Resolution variation maps for the final 3D reconstruction. **f**. Gold standard FSC plots for the final 3D reconstructions for the two maps. **g**. Plots of the particle number vs the reciprocal squared resolution. The B-factor was calculated as 2⨯the linear fitting slope.

**Figure S2.**
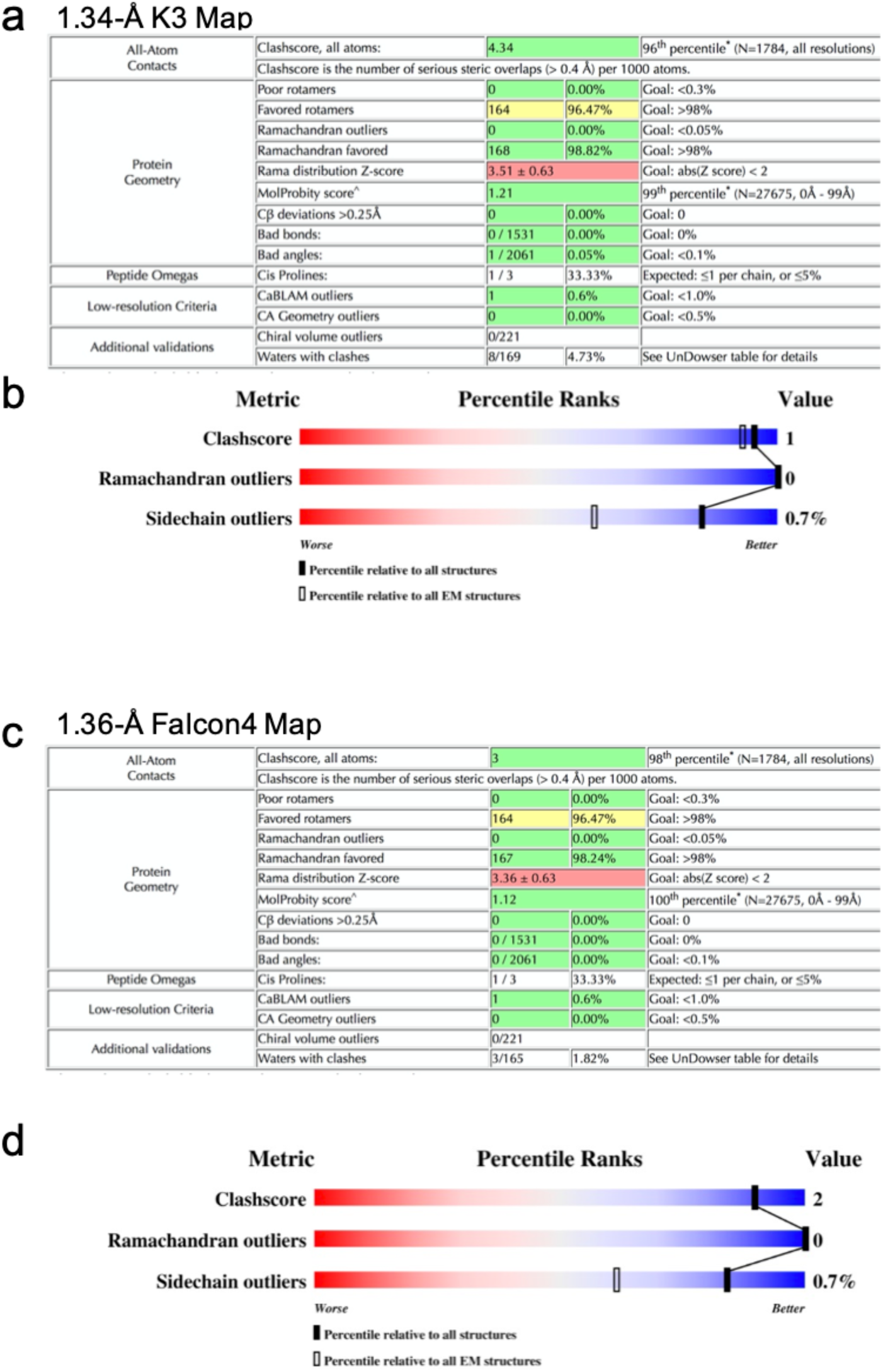
Assessment of model quality. MolProbity analysis (a, c) and overall quality derived from PDB validation reports (b,d) reports for atomic models fitted to 1.34 Å (a, b) and 1.36 Å (c, d) maps. Both reports show good model geometries and very few water atoms that clash with other atoms.

**Figure S3.**
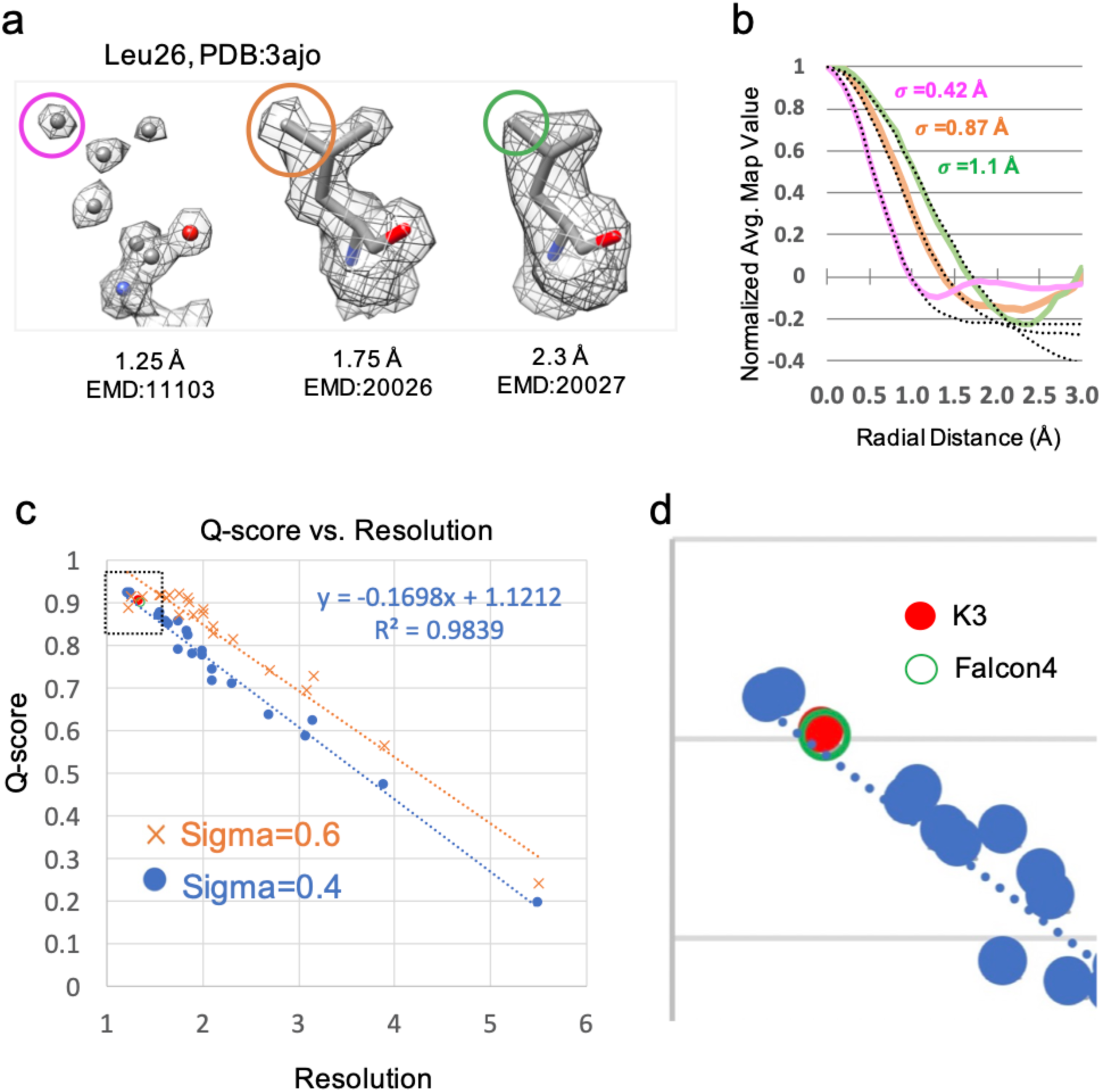
Q-score adjustment for apoferritin maps at better than 1.5 Å resolution. Atoms in maps at 1.25 Å, 1.75 Å, and 2.3 Å resolution (a) all have very close to Gaussian-like atomic profiles with smaller widths (σ) at higher resolutions (b). Previously, Q-scores were calculated using a sigma parameter of 0.6, which gave the highest Q-score of 1 at a resolution of ∼1.5 Å. With this parameter, Q-scores start to drop at higher resolutions as shown in (c). Thus, we adjusted the Q-score calculation to give the highest score of 1 at ∼1.2 Å by setting sigma to 0.4; with this parameter, the plot of Q-score vs. resolution for 10 maps & models in the EMDB is now linear again. Panel (d) shows a close-up of (c) at resolutions between 1-2 Å, highlighting the Q-scores for the new maps of apoferritin at 1.34 Å (K3) and 1.36 Å (Falcon 4).

**Table S1.** Cryo-EM data collection and processing

| Apoferritin | K3 dataset | Falcon4 dataset |
| --- | --- | --- |
| **Data collection and processing** |  |  |
| Microscope | Titan Krios G3i | Titan Krios G3i |
| Voltage (kV) | 300 | 300 |
| Camera | Gatan K3 | Thermo Fisher Falcon4 |
| Grids Type | R2/1 Quantifoil grids | R2/1 Quantifoil grids coated with continuous carbon film |
| Sample concentration | 1.5 mg/ml | 0.2 mg/ml |
| Magnification | 215,000x | 155,000x |
| Pixel size (Å) | 0.4 | 0.502 |
| Total exposure (e-/Å<sup>2</sup>) | 45 | 40 |
| Exposure time (s) | 0.5 | 2 |
| Number of frames per exposure | 30 | 40 |
| Energy filter slit width (keV) | 15 | N/A |
| Data collection software | EPU | EPU |
| Defocus range (µm) | -0.35 - -1.3 | -0.3 - -1.3 |
| Number of micrographs collected | 8,034 | 7,734 |
| Number of micrographs used | 6,951 | 5,427 |
| Number of initial particles | 1,176,336 | 707,350 |
| Symmetry | O | O |
| Number of final particles | 902,455 | 500,643 |
| Resolution (0.143 gold standard FSC, Å) | 1.34 | 1.36 |
| Local resolution range (Å) | 1.3 – 1.5 | 1.3 – 1.5 |

